# Face recognition depends on specialized mechanisms tuned to view-invariant facial features: Insights from deep neural networks optimized for face or object recognition

**DOI:** 10.1101/2020.01.01.890277

**Authors:** Naphtali Abudarham, Galit Yovel

## Abstract

Faces are processed by specialized neural mechanisms in high-level visual cortex. How does this divergence to a face-specific and an object-general system contribute to face recognition? Recent advances in machine face recognition together with our understanding of how humans recognize faces enable us to address this fundamental question. We hypothesized that face recognition depends on a face-selective system that is tuned to view-invariant facial features, which cannot be accomplished by an object-general system. To test this hypothesis, we used deep convolutional neural networks (DCNNs) that were optimized for face or object recognition. Consistent with the hierarchical architecture of the visual system, results show that a human-like, view-invariant face representation emerges only at higher layers of a face-trained but not the object-trained neural network. Thus, by combining human psychophysics and computational modelling of the visual system, we revealed how the functional organization of the visual cortex may contribute to recognition.

---

It is well established that faces are processed by a specialized neural mechanism in high-level visual cortex, which is highly selective to faces relative to all other object categories (1–3). However, it is still unclear how this functional specialization may contribute to face recognition. Face recognition is a computationally challenging task that requires generalization across different appearances of the same identity, as well as discrimination between different identities, for a large set of homogeneous visual images. In order to perform this taxing task, a system needs to learn which features are useful for both discrimination across identities as well as for generalization across different appearances and views of the same identity. Here we hypothesize that view-invariant face recognition depends on neurons that are specifically tuned to view-invariant facial features, whereas an object recognition system is tuned to features that do not support view-invariant face recognition. Such an account would underscore the need for a dedicated face-selective mechanism for face recognition.

Recent developments in machine face recognition, enable us to test this hypothesis, with deep convolutional neural networks (DCNNs). In particular, face and object-trained DCNNs have recently reached and even surpassed human face and object recognition abilities (4, 5). This human-level performance together with their brain-inspired hierarchical structure, make them good candidates to model the human brain (6). There are two main advantages for using DCNNs as models of human visual processing: first, we can fully control the type of stimuli that these models are trained with, and this way separately model a face or an object recognition systems by training with only faces or only objects. Second, examination of the representations that are generated across the layers of DCNNs have shown that earlier layers represent low-level visual features, similar to primates’ early visual cortex, whereas high-level layers represent high-level visual features that support object/face recognition (7). To this end, we used a DCNN that is trained to recognize objects, to model an object-processing system, and a DCNN that is trained to recognize faces as a model for a face-processing system. We thereafter compared the representations that these two DCNNs generate for face images across their different processing layers, and tested whether and at what stage during the processing hierarchy they generate a view-invariant representation of face identity. Based on the structure of the primate visual system, where the division to face and object areas emerges at higher levels of the visual system, we expect that higher-layers of the face-trained, but not object-trained DCNN, will generate a human-like view invariant face representation. In contrast, lower-level layers of both networks will generate similar, image-based face representations.

In order to examine whether a DCNN generated a human-like, view-invariant representation across its different layers, we used two measures of human view-invariant face recognition: First, we examined how similar are the representations of face identity across different head views (Study 1). Second, we examined the sensitivity of the DCNNs to view-invariant facial features that are used by humans for face recognition (8–10). For both measures, which will be described in more details below, we examined the representation of DCNNs trained to recognize faces or objects, across their layers, and compared them to the representation generated by humans.

## Study 1

In order to examine whether the representation of face identity is view invariant or image-specific, we obtained two types of measures: First, we measured the distances between representations generated for images of the same identity in different head views. A view-invariant representation is indicated by smaller distances for same-identity images across different views, compared with images of different identities from the same view (Figure 2). Second, we computed the distances between pairs of faces of different identities that were in the same head-view, and correlated these distances with distances for the same pairs across the different views (Figure 3A) (similar to Representation Similarity Analysis (11)). High correlations across head views, indicate a view-invariant representation of face identity, whereas low correlations reflect an image-specific representation. This measure was also used to quantify human view-invariant face representation.

### Results

#### DCNN performance on face and object verification tasks

Figure 1B shows the performance of the face and object-trained networks on face and object verification tasks. Performance for faces was better than objects in the face-trained DCNN, whereas performance for objects was better than faces in the object-trained DCNN. Performance of the face-trained DCNN (96.5%) was close to human level performance (97.53%) on the LFW benchmark (http://vis-www.cs.umass.edu/lfw/results.html#Human). This performance is somewhat lower than the current state-of-the-art (http://vis-www.cs.umass.edu/lfw/results.html) because the training set that we used was limited to 300K faces (see Methods section for details) but is still very close to human performance.

**Figure 1:**
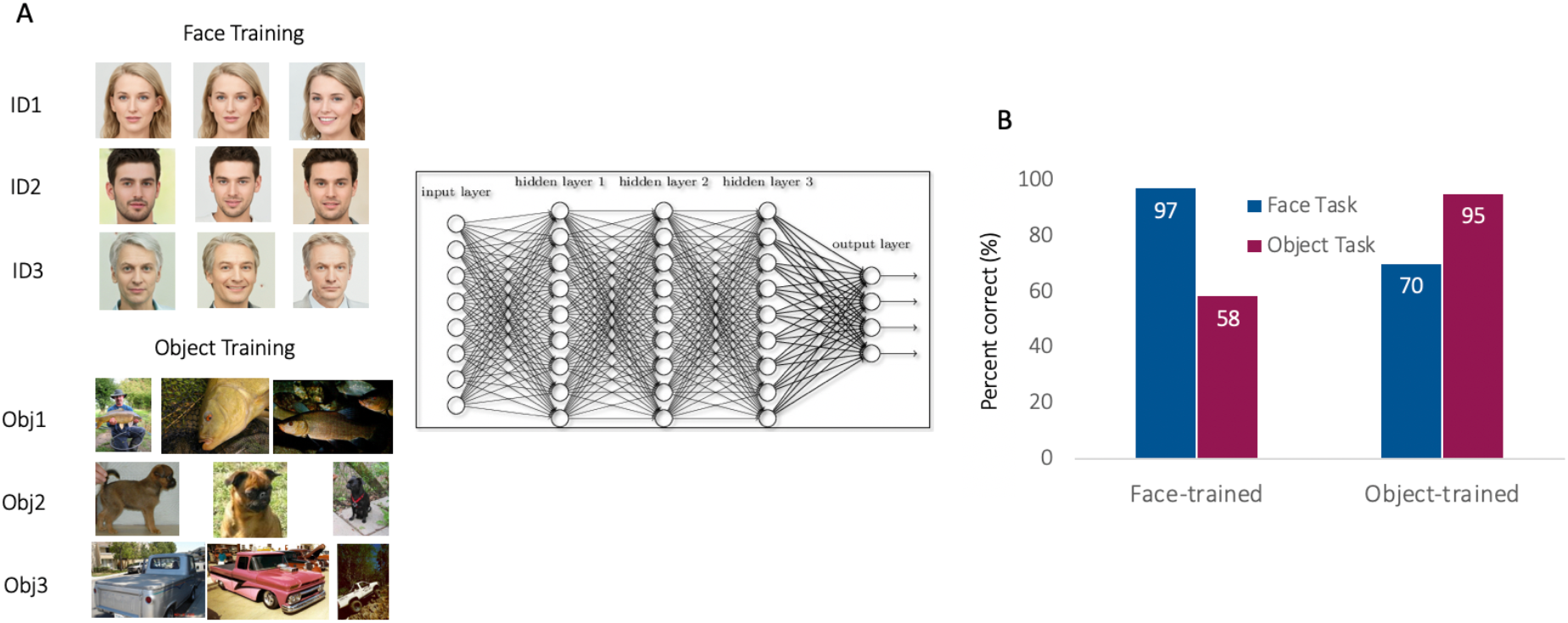
A. A DCNN was trained to recognize 1000 object categories or 1000 face identities. B. Face verification performance of the two networks was evaluated with the standard Labeled Faces in the Wild (LFW) benchmark for faces, and a similar verification task with untrained object images. **Face images were replaced by AI-generated faces in this manuscript due to bioRxiv’s policy on not including real human faces within posted manuscripts. The experiment stimuli included real human photos from the VGGface database.**

#### The representation of identity across head-views

As explained above, we used two measures to quantify whether the representation of face identity is view-invariant:

##### Distances between same identity across head-views relative to different identity

Figure 2 depicts the average normalized Euclidian distances between the representations generated by the face-trained and object-trained DCNNs, for pairs of images of the 3 same-identity conditions – same identity frontal view, same identity quarter-left view, same identity half-left view and the different identity-same frontal view (see Figure 2A) in the last layer (Figure 2B) and across all layers (Figure 2C).

**Figure 2:**
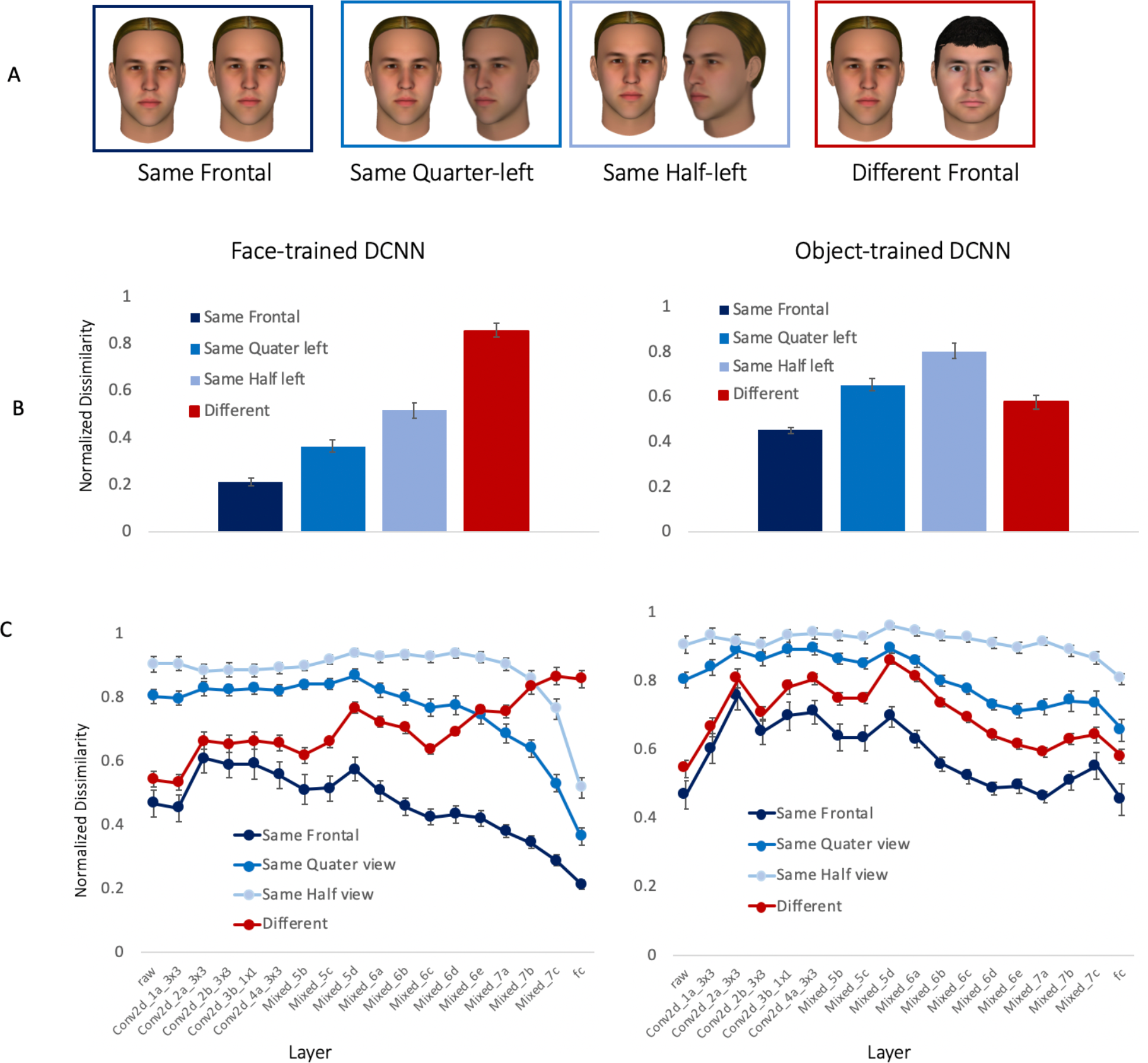
A. To quantify view invariance, pairs of same identity faces from the same frontal view or different views as well as same-view different identity faces were used. B. The normalized Euclidian distances between the representations of each pair were computed for the last layer of the face-trained and object-trained DCNNs. Results show higher dissimilarity for different identity same-view faces than same identity faces across head views – indicating an identity-specific view-invariant representation in the face-trained DCNNs. The object-trained DCNN shows higher dissimilarity for same identity different-view faces, than for different identity same-view faces, indicating a view-specific representation. C. The representations across the layers indicate a view-specific representation in both the face and object-trained networks for low-level and mid-level layers, but a view-invariant representation only for the higher-layers of the face-trained network. **Face images were replaced by illustrations in this manuscript due to bioRxiv’s policy on not including human faces within posted manuscripts. The experiment stimuli included real human photos**

A repeated measure ANOVA with DCNN training (Face, Object) and Head-view (Same Frontal, Same quarter left, Same half left and Different frontal) on distances between face pairs in the last layer of the two DCNNs, revealed a significant interaction (F(3,24) = 45.14, p <.0001, η^2^_p_ =.85). This interaction reflects the view-invariant representation in the face-trained and view-specific representation in the object-trained networks (Figure 2B). In the face-trained DCNN, the distances between different identity faces was the largest and significantly different from distances between same identity faces across the different head views (p <.001, corrected for multiple comparisons, Cohen’s d = 2.5 – 4.46). In contrast, for the object-trained DCNN the distances between different identity faces were significantly smaller than the distances for same identity different-view faces (p <.005, Cohen’s d = (−1.9) – (−4.3)). In addition, there was no significant difference between same and different frontal faces (p =.65, corrected for multiple comparisons, Cohen’s d =.079).

We also examined the representations across the different layers (Figure 2C). A significant interaction between DCNN training, Head-view and Layer (F(48,384) = 37.3, p <.0001, η^2^_p_ =.82), indicates that overall, the representations in the different DCNNs were similar for the initial layers, which showed a view-specific representation, but different for higher layers, which remained view-specific for the object-trained DCNN but became view-invariant for the face-trained DCNN. This was indicated by higher dissimilarity for same identity faces across head views than different identity and same identity faces within the same head-view in lower layers of the face and object-trained DCNNs. In the higher layers of the DCNNs we see a view invariant representation of face identity for the face-trained but not the object-trained DCNN.

Next, we compared the DCNN representations to human representations. In principle, we could have collected human’s similarity judgements on pairs of faces and compare to the DCNN distance measures reported above. However, since humans can easily generalize across different views of the same identity, they would give the same similarity measures across different head-views. We therefore used an indirect measure of view-invariance in humans and compared that to a similar measure in DCNNs.

##### Correlations between similarity scores across head-view

To obtain an indirect measure of view-invariance, we computed the correlations between similarity distances within the same head view conditions (for details see Methods section) (Figure 3A). Briefly, we correlated distances between each pair of different identities across the different head views. High correlations indicate a view invariant representation (i.e. the distances btw face1-face2, face2-face3 etc. are correlated across head views). Low correlations indicate a view-specific representation.

**Figure 3:**
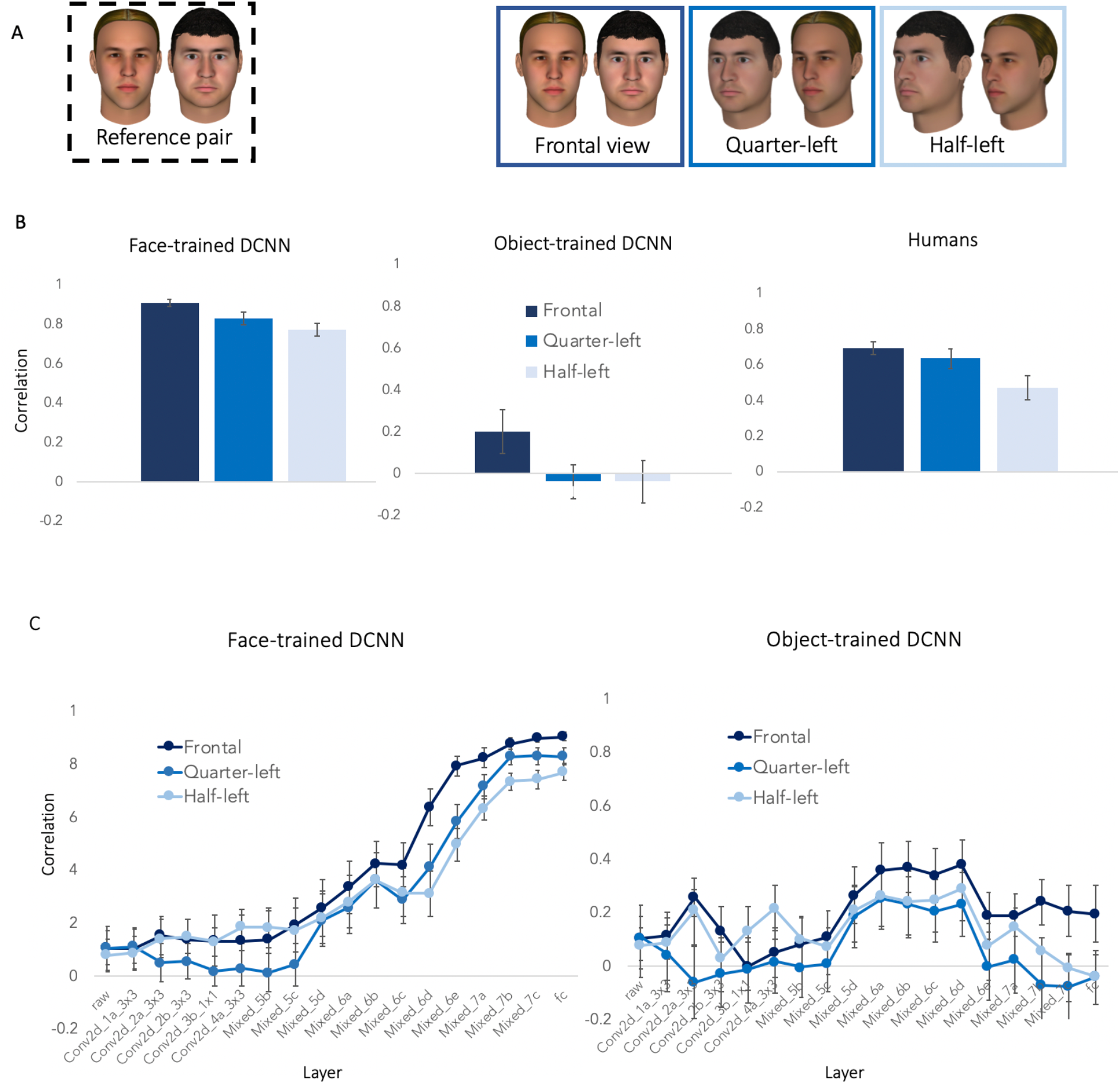
A. Distances between pairs of different identities, for each of the different views, were correlated with the distances between pairs of faces in a reference set of frontal faces. B. Correlations across head views for the last layer of the face-trained, object-trained and human similarity measures. High correlations indicate a view invariant representation of face images, and low correlations – an image-specific face representation. Correlations across head-views were higher for frontal than quarter left and half left views in the face-trained DCNN and in humans. For the object-trained DCNNs correlations were very low indicating an image specific representation. C. Examination of the representations across DCNN layers indicated an image-specific representation in lower-layers of the face and object-trained DCNNs, and a view-invariant representation of faces in higher layers of the face-trained DCNNs. **Face images were replaced by illustrations in this manuscript due to bioRxiv’s policy on not including human faces within posted manuscripts. The experiment stimuli included real human photos**

For the face-trained DCNN, the correlations were overall high with a monotonic decrease for larger head-view differences. In contrast, for the object-trained DCNN the correlations were overall low, indicating an image-based representation (Figure 3B).

To compare the representation of the DCNNs to humans, we computed the same correlations for humans (see methods). Each participant was presented with only one head-view so they see each identity pair only once, and correlations were performed for measures obtained from different groups of participants. Human results revealed a pattern that was similar to the face-trained DCNN. A repeated measure ANOVA with System (Human, Face-trained DCNN, Object-trained DCNN) and Head-view (Frontal, Quarter-left, Half-left) revealed a main effect (F(2,18) = 81.09, p <.001, η^2^_p_ =.90), indicating a much higher correlation for the human and face-trained than the object-trained DCNN. In addition, the correlations of the face-trained DCNN were higher than humans (F(1,10) = 14.50, p <.001, η^2^_p_ =.59) indicating a better view invariant representation in face-trained DCNN than humans. It is noteworthy that human similarity scores were correlated between different groups of participants, which might have lowered the overall level of correlations. Still correlations were much higher in humans than the object-trained DCNN. There was no interaction between System and Head-View indicating a similar pattern of higher correlation for the frontal than quarter-left and half-left faces for humans and DCNNs (F(4,36) = 0.07, p =.77, η^2^_p_ =.05).

We also examined the correlations across the different layers of the face-trained and object-trained DCNNs (Figure 3C). A repeated measure ANOVA with Training (Face, Object) × Head-View (frontal, quarter, half) and Layer revealed a significant interaction between Train and Layer (F(16,160) = 79.93, p <.0001, η^2^_p_ =.89). This interaction reflects low correlations across head-views, indicating an image-based representation, for the face and object-trained DCNNs at lower-layers. Whereas at higher layers of the DCNNs correlations were much higher in the face-trained but not the object-trained DCNN. These findings indicate a similar image-specific representation in lower-layers of the face and object-trained DCNNs, and a view-invariant representation at higher levels of processing for the face-trained but not the object trained DCNNs. These results are in line with the distance measures we reported in the previous section.

### Discussion

Results of Study 1 show that a human-like view-invariant representation of face identity emerges at higher levels of processing of a system that is specifically tuned to faces. The face-trained DCNN was trained on different appearances of many different identities and this way learned which features are useful for the generation of a view-invariant face representation. This representation was similar to the representation generated by humans using the same measure to quantify view-invariance. Importantly, a network that was trained with objects and was able to discriminate many categories of objects across their different appearances, did not generate a view-invariant representation of face identity. The representation of the object-trained network was view-specific and was similar across its different layers to the representation that was generated in lower-level layers of the face-trained network.

The emergence of a view-invariant representation at higher-level of face processing, following a view-specific representation at lower and mid-level face areas, is in line with findings reported in single unit recordings of face neurons in different face patches along the hierarchy of the face network of the macaque (12). In particular, a view-specific representation that was not sensitive to identity was found in the posterior face-area (area ML) whereas a view-invariant, identity-selective representation was found in the more anterior face-area (area AM). This pattern of response parallels the DCNN representations of the face-trained but not the object-trained network, and indicates that the view-invariant representation depends on a system that is tuned to view-invariant, high-level facial features. To further examine this suggestion, in Study 2 we tested whether the face-trained but not the object-trained system is tuned to view-invariant facial features that are used by humans for face recognition.

## Study 2

In a series of previous studies, we discovered a subset of view-invariant facial features that are used by humans to recognize faces (9, 10, 13). This subset of facial features includes the hair, lip-thickness, eye-color and shape and eyebrow-thickness. Our findings showed that humans can reliably discriminate between these features across head views of different identities, indicating that these features are useful for both generalization and discrimination (13). We further found that when these features are modified, faces cannot be identified and are judged as different identities (see Figure 4A, for an example of George Clooney). We therefore named these view-invariant features *critical features*, as they are critical for the identity of a face. In contrast, another set of features including eye-distance, face-proportion, mouth-size or nose-shape were not reliably discriminated across head-views of different identities, and changing faces by modifying these view-dependent features did not change the identity of a face. These features were therefore named *non-critical* features.

**Figure 4:**
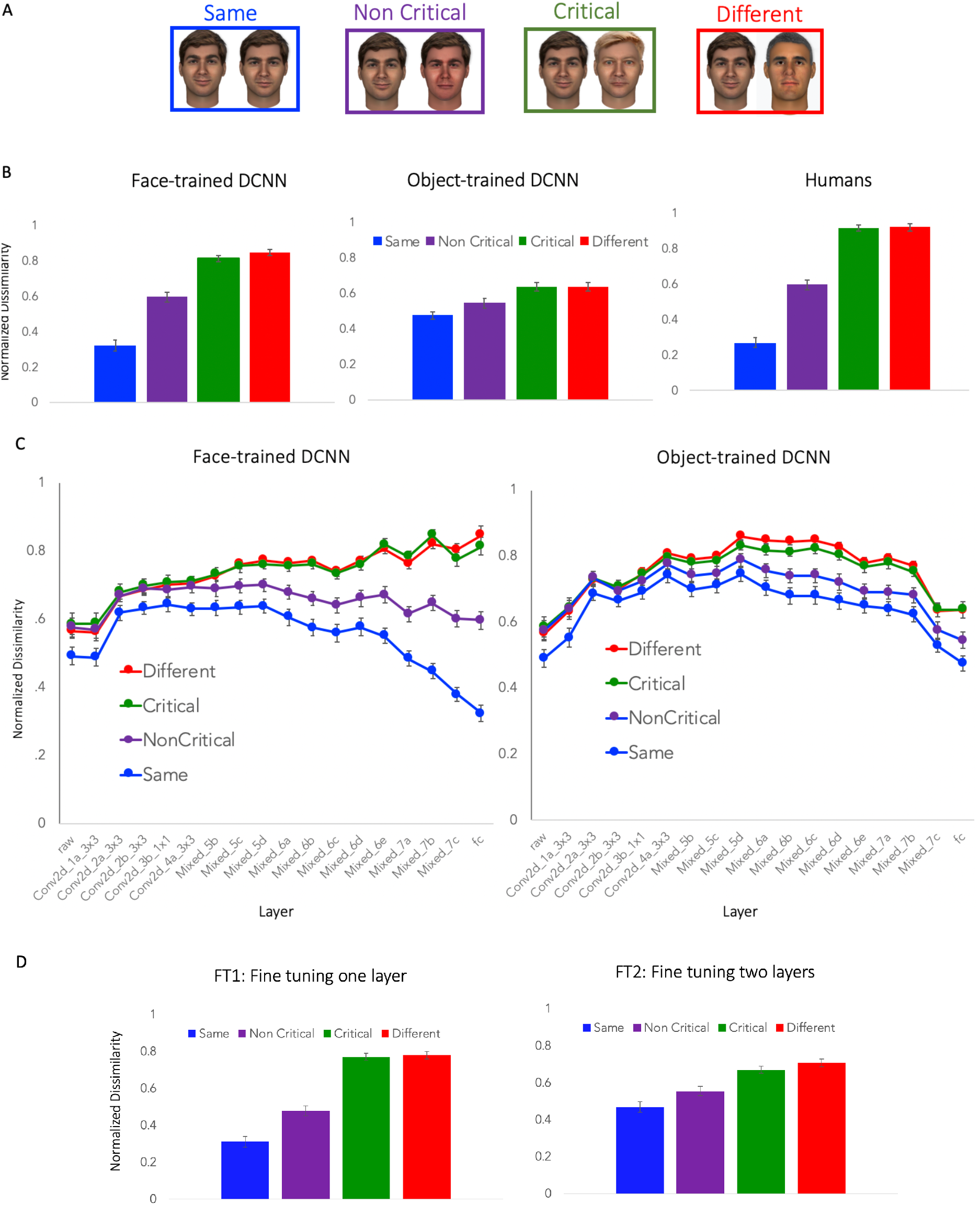
A. An example of the four conditions with the face of George Clooney, including a pair of same identity faces, a pair in which non-critical features were modified, a pair in which critical features were modified and different identity faces. B. Distances between the representations of the four different pairs in the last layer of a face-trained DCNN, object-trained DCNN, and similarity measures obtained from humans, for the same face images, indicate higher sensitivity to critical features in humans and the face-trained DCNN relative to the object-trained DCNN. C. The representations across the different layers show that sensitivity to critical /view-invariant features emerges at higher layers of the face-trained DCNN. Low level layers of both DCNNs and high-level layers of the object-trained DCNN were less sensitive to critical features as well as to face identity. D. Training of the last layers of the object-trained DCNN with faces (fine-tuning) indicate that fine tuning of the last 2 trainable layers generated a representation that was similar to the fully trained face DCNN. **Face images were replaced by illustrations in this manuscript due to bioRxiv’s policy on not including human faces within posted manuscripts. The experiment stimuli included real human photos**

We hypothesized that these critical features are learned through our experience with different identities as well as different appearances of each of these identities, to enable generalization across different views of the same identity, as well as discrimination between different identities. Based on results of Study 1, if these features support view-invariant representation of face identity, we expect that a face-trained DCNN will be sensitive to these features, in particular in its higher layers, whereas an object-trained DCNNs will be less sensitive to these features. We also expect that the face and object-trained DCNNs will be both insensitive to these features in lower-level layers of the networks.

To examine the sensitivity of the DCNNs to these features, we measured the distances between face-representations of two types of face-pairs: an original face vs. the same identity in which we modified critical features, and an original face vs. the same identity in which we modified non-critical features (see Figure 4A). If the face-trained DCNN uses the same critical/view-invariant features as humans, then the distances between faces that differ in critical features will be larger than faces that differ in non-critical features. In addition, as a baseline, we also measured the distances between unchanged images of same identities, and the distances between images of different identities.

### Results

#### The Representation of View-invariant Features in Face-trained and Object-trained DCNNs

We measured the Euclidean distances between the representations of face images in the 4 conditions across all layers of the face-trained and object-trained DCNNs. Fig. 4B shows the comparison of the average distances across all face identities calculated using the last layer of the face-trained and the object-trained DCNNs, as well as similarity distances made by humans (adapted from (10)). A repeated measure ANOVA with Training Type (Face, Object) and Face Type (Same, Non-Critical, Critical, Different) on dissimilarity scores of 25 face identities revealed a significant interaction between the two factors (F(3,72) = 70.19, p <.001, η^2^_p_ =.75). Planned comparisons show that the difference between critical and non-critical features was significantly larger in the face-trained (Cohen’s d = 1.56) than the object-trained DCNN (Cohen’s d = 0.67) (F(1,24) = 13.31, p <.001, η^2^_p_ =.36).

We then compared the representation of the last layer of each of the DCNNs to human’s similarity ratings of the same stimuli. A mixed ANOVA with System (Human, DCNN) as a between-subject factor and Face Type (Same, Non-Critical, Critical, Different) on dissimilarity scores of 25 face identities revealed a significant interaction for humans and object-trained DCNN (F(3,144) = 135.58, p <.001, η^2^_p_ =.74) and a much smaller difference between humans and face-trained DCNN (F(3,144) = 9.23, p <.001, η^2^_p_ =.16). As can be seen in Figure 4B, distances between the different pairs were relatively similar for the object trained-network, whereas both the human and the face-trained DCNN showed a larger difference between same and different identity pairs, as well as a larger difference between critical and non-critical feature changes.

We then examined the sensitivity to critical features across the different layers of the face-trained and object-trained DCNNs. Inspection of Figure 4C shows that the sensitivity to face identity in general (difference between Same and Different pairs) and to critical features in particular (difference between critical and non-critical features), emerges at higher layers of the face-trained DCNN. The representation in earlier layers of both DCNNs and the higher layers of the object-trained DCNN was less sensitive to face identity as well as to critical features. Indeed, a repeated measure ANOVA with Training Type (Face, Object), Layer (all 17 layers) and Face type (Same, Non-Critical, Critical, Different) on the dissimilarity scores of 25 face identities, revealed a significant interaction of the three factors (F(3,1152) = 43.65, p <.001, η^2^_p_ =.65). The effect of Face Type indicates two important findings: First, the difference between same and different identity pairs was smaller in early layers of both networks, as well as in the higher layers of the object-trained DCNN, whereas the difference was very large in the last layers of the face-trained DCNN, indicating discrimination across different identities. Similarly, the difference between faces that differ in critical and non-critical features is largest in last layer of the face-trained DCNN, and smaller in the last layer of the object-trained DCNN and early layers of both networks. Indeed, the correlations between distances of Different-Same pairs and distances between Critical – Non Critical pairs across the different layers was 0.96, indicating that higher sensitivity to critical features is associated with better discrimination of face identity.

#### Face training for specific layers (i.e. Fine Tuning) of the object trained DCNN

Comparison of the face-trained and object-trained DCNNs indicates that the representation of critical and non-critical images at early layers of networks is similar. These findings indicate that selective training of the final layers of the object-trained network on face recognition may suffice to generate a view-invariant representation of face identity.

As can be seen in Fig. 4C, the stage in the hierarchy of processing where the representations of the two DCNNs diverge is at 3-5 last layers of the DCNN. To further examine the exact point of divergence we took the object-trained DCNN, and trained only the last layers on the dataset that was used to train the face-trained DCNN (a procedure known as “fine-tuning”), using the same training procedure. We then examined how many layers of the object-trained DCNN should be trained with faces to obtain the same representation that a DCNN that is fully-trained with faces generates (see Figure 4D).

First, we trained only the last trainable layer (Layer “Mixed_7c” see methods for information). We name this condition fine-tuning 1 (FT1). A repeated measure ANOVA with Training type (Full, FT1) and Face Type (Same, Non-Critical, Critical, Different) on the representation of the last layer revealed a significant 3-way interaction (F(3,72) = 29.88, p <.001, η^2^_p_ =.55), indicating highly different face representations for the two DCNNs. Next, we repeated the same procedure, only now we trained the last 2 trainable layers of the DCNN (i.e. “Mixed_7c” and “Mixed_7b”). We name this condition fine-tuning 2 (FT2). Here, the 3-way interaction was not significant (F(3,72) = 1.05, p =.38, η^2^_p_ =.04). As can be seen in Figure 4D, the representations of a fully-trained DCNN and a DCNN in which only the last 2 layers were trained with faces are nearly identical (Figure 4D).

Finally, to assess whether sensitivity to critical features is associated with better performance on a face recognition task, we measured the performance of each of the layers of the face-trained DCNN on the benchmark face (LFW) verification task, as explained in the Results section of Study 1. We then computed a measure of sensitivity to critical features by subtracting the distances between the representations of pairs that differ in critical features and pairs that differ in non-critical features for each layer. A higher difference indicates greater sensitivity of the layer to critical features. We then computed the correlations between the two measures. Figure 5 shows a very strong correlation (r = 0.98) indicating that increased sensitivity to critical features used by humans, is associated with improved performance of the DCNN on a benchmark face recognition task (LFW).

**Figure 5:**
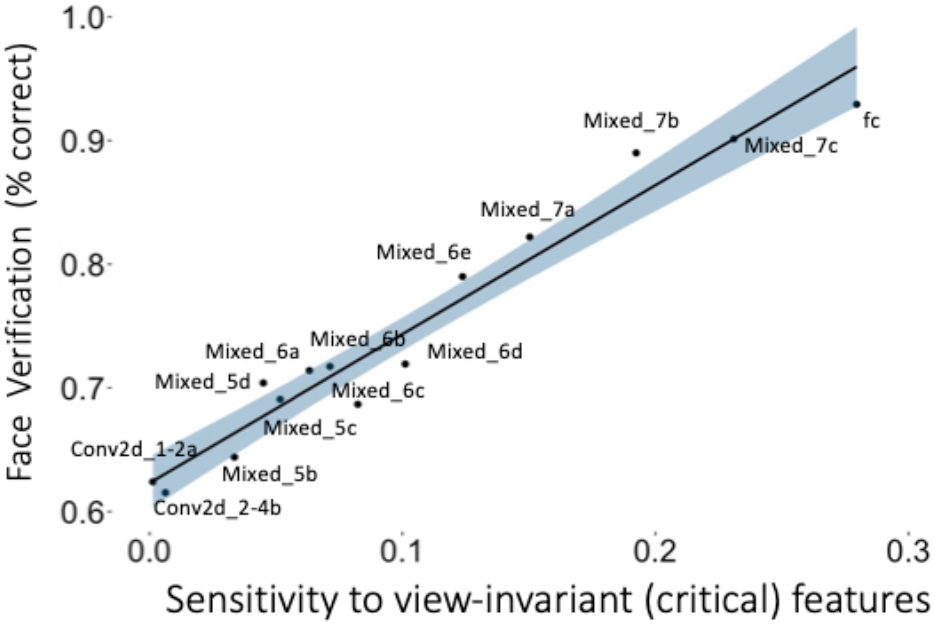
Performance on a benchmark face verification task (LFW) for each of the 17 layers of the network is highly correlated with the sensitivity of each layer to view invariant/critical features. Higher face recognition ability is associated with higher sensitivity to critical features.

### Discussion

Results of Study 2 indicate that the face-trained but not the object-trained DCNN uses the same critical/view-invariant features used by humans for face recognition. In addition, the sensitivity to these view-invariant facial features emerges only at the higher layers of the face-trained DCNN, whereas low-level layers represent low-level features that are useful for both face and object recognition. Indeed, retraining of the last two layers of an object-trained network with faces, generated a similar representation and performance level on a face verification task as a DCNN that was fully trained with faces. Finally, the sensitivity to these view-invariant critical features was strongly correlated with DCNNs performance on a face verification task (Fig. 5). Higher layers showed increased sensitivity to critical features than lower layers and also improved performance on a benchmark face verification task (LFW).

These findings are in line with results of Study 1 that showed that a view-invariant representation emerges only at the higher layers of a face-trained network. We therefore suggest that both humans and a face-trained DCNN relay on the same high-level, view-invariant features to enable view-invariant face recognition. It is important to note that we do not suggest that face-trained DCNNs are specifically tuned to measure lip-thickness or eye-shape, nor do we suggest that human brain neurons are tuned that way. We do suggest, however, that the type of information that human neurons and DCNN units rely on for face processing is correlated with the view-invariant features that we manipulated. Taken together, we conclude that view-invariant face recognition depends on a system that is specifically tuned to high-level, view-invariant facial features.

### General Discussion

The question of why faces are processed by a specialized mechanism has been extensively discussed in the face processing literature (e.g., (1, 2)). By using face-trained and object-trained DCNNs as models for this divergence to a face processing and an object processing systems, we show that face recognition depends on specialized mechanisms that are tuned to high-level, view-invariant facial features, which cannot be achieved by a system that is tuned to features that support object recognition. Our findings also show that at early stages of processing, faces and objects rely on the same visual information. It is only at the final, higher-level stages, that the two systems diverge to a specialized system that is finely tuned to facial features, and to a general object system that processes faces and objects similarly, but cannot support face recognition, similar to the primate visual system. A similar divergence at higher layers of processing was recently discovered for auditory stimuli between processing of speech and music in a task-optimized DCNN that generated human-like performance (14).

A prevalent alternative account for the face-specific hypothesis, is that the specialized mechanism for face processing supports recognition of any non-face object of expertise, that involves discrimination at the individual level. Indeed, a basic difference between object and face recognition is that the former is categorized at the basic level (e.g. a car vs. a truck), whereas faces are categorized at the individual level (Joe vs. Danny). It has been therefore suggested that the face system is not specialized for faces but for discrimination of any category at the individual level (e.g. specific cars or birds) (15, 16). The current findings, which indicate that face processing relies on face-specific features, implies that expertise with other categories is likely to generate a separate system that is specialized for features needed for the object of expertise (e.g., (1, 17)). The computational approach used here can be also used to test the predictions of the expertise hypothesis and the face-specific hypothesis, by studying categorization of individual members of non-face categories in face-trained and object-trained networks. For example, we can examine whether re-training with non-face objects at the individual level lead to better performance in a face-trained than an object-trained DCNN, making the face-specific system more suitable for any type of individual based categorization.

Our findings highlight the importance of specific training with faces for the generation of a view-invariant facial features. Such training enables the system to learn which features are both invariant across different appearances of the same identity and are also useful for discrimination between identities. The features that we tested here are based on results of our previous studies that used faces of adult male Caucasian faces, and may not generalize to faces of other races. For example, hair and eye color, which are both invariant and discriminative for Caucasian faces, are invariant in Asian and African faces but may not be discriminative for these races. Indeed, it is well established that humans show poor recognition for races for which they have low experience with, an effect known as the Other Race Effect (e.g., 13). Similarly, DCNNs were shown to be biased for the races that are included in their training set. State of the art and commercial algorithms show much lower performance for African and Asian faces than Caucasian faces (19) Thus, both human and DCNN representations indicate that the features that the face system is tuned to may not be selective only to faces, but to facial features that are useful for a specific category of faces we have experience with. This further highlights the degree of specificity in visual experience that is required for intact face recognition.

Finally, DCNNs have been criticized for being “black box” machines that have millions of parameters, and therefore reverse engineering their underlying feature representation is a great challenge (6, 20). Here we show that insights from reverse engineering of the human mind, and the discovery of features that are used by humans, can shed some light on the type of information used by DCNNs to accomplish their human-level performance. Whereas a possible disadvantage of the features used in our studies, relative to other data-driven reverse-engineering techniques (e.g. bubbles (21); layer-visualization; layer activations (22)), is that they are limited to features that have semantic meaning, their advantage is that they can specify different features in the same spatial location that may be useful or not useful for face recognition, whereas data-driven visualization techniques highlight specific spatial locations in the image that are diagnostic for recognition. For example, lip-thickness was found to be critical for face recognition, but mouth-size found in the same spatial location was not. This enables us to refine our tests for feature sensitivity and reveal a remarkable similarity between the representations used by humans and face-trained DCNNs for face recognition. This similarity between the perceptual representations of humans and machines is not trivial given the many differences between the architecture and computations performed by the human brain and a feed-forward DCNN. However, these findings indicate that feed-forward DCNNs can make good models of early to mid-level stages of visual processing. A similar approach can be used with any category as well as in other modalities (e.g. voices) to reveal its critical features in humans and examine whether it can also account for machine performance.

In summary, we found that a face-trained, but not an object-trained DCNN, generates a human-like view-invariant face representation. Furthermore, this representation emerges at higher layers of the face-trained network, while lower layers of both networks generated a similar view-specific representation. These finding may resemble the neural pathways for processing faces and objects, that are similar in the low-level visual areas, and diverge to dedicated modules in higher levels of the visual system. We suggest that humans and face-trained DCNNs learn to use the same invariant features based on experience with faces in variable views, and that this view-invariant representation cannot be learned from experience with non-face objects. We therefore conclude that human-like view-invariant face recognition cannot be accomplished by an object-general system but depends on a specialized face system that is tuned to high-level, view-invariant facial features.

## Methods

### Study 1

#### Stimuli

To quantify view-invariance we used images of 15 identities from the color FERET face-image dataset (23, 24). For each identity we took 4 images: 1 – a frontal image, hereby referred to as the “reference” image, 2 – a second frontal image, different from the “reference” image, hereby referred to as the “frontal” image, 3 – a quarter-left image, and 4 – a half-left image (see Figure 2A – for example images). All face images were of adult Caucasian males, had adequate lighting, with no glasses, hats or facial hair. The images were cropped just below the chin to leave only the face, including the hair and ears.

#### Face-trained and Object-trained DCNNs

For the Object-trained DCNN we used the pre-trained inception_v3 DCNN from https://pytorch.org/docs/stable/torchvision/models.html, that was trained to recognize the 1000 categories of the ImageNet Large Scale Visual Recognition Challenge (ILSVRC, http://image-net.org/challenges/LSVRC/). For face training we took the same inception_v3 DCNN (defined in https://github.com/pytorch/vision/blob/master/torchvision/models/inception.py), and trained it on a subset of the VGGFace2 face image dataset (http://www.robots.ox.ac.uk/~vgg/data/vgg_face2/), by selecting the first 1000 identities from the dataset, and the first ~300 images for each identity as train data, and ~50 images per identity as validation data. We started with random weights, using the default training parameters of https://github.com/pytorch/examples/blob/master/imagenet/main.py and trained the network for 120 epochs of 1000 iterations each.

To measure the DCNN level of performance on a face verification task, we used the standard Labeled Faces in the Wild (LFW) benchmark (http://vis-www.cs.umass.edu/lfw/). To measure the DCNN level of performance on an object verification task, we constructed a similar task by randomly selecting 4000 untrained image-pairs from the validation set of the ILSVCR2012 dataset (2000 were same-category image-pairs and 2000 were different-category image-pairs).

#### Extracting representations from DCNNs

To extract the representations that were generated by the DCNNs, we ran the trained models in evaluation mode on a predefined set of image stimuli (see Stimuli section above). The face images were first aligned using the python API of the “dlib” library (http://dlib.net/python/index.html), with the face landmark predictor http://dlib.net/files/shape_predictor_68_face_landmarks.dat.bz2. This library failed to detect 4 of the faces in the half-left angle, so they were removed from analysis. Following alignment, the images were normalized with the standard ImageNet normalization (mean=[0.485,0.456,0.406], std=[0.229,0.224,0.225]).

We first examined the last pooling layer of the network, the input to the fully-connected layer of the network (the input to layer “fc” in https://github.com/pytorch/vision/blob/master/torchvision/models/inception.py). This is the layer that generates the final representation (last layer) that is later transformed to the output layer, which estimates category probabilities. In Figures 2–4 we refer to it as “fc”. We then examined the representations across the different layers of the network. For that we used layers: “Conv2d_1a_3×3”, “Conv2d_2a_3×3”, “Conv2d_2b_3×3”, “Conv2d_3b_1×1”, “Conv2d_4a_3×3”, “Mixed_5b”, “Mixed_5c”, “Mixed_5d”, “Mixed_6a”, “Mixed_6b”, “Mixed_6c”, “Mixed_6d”, “Mixed_6e”, “Mixed_7a”, “Mixed_7b”, “Mixed_7c”.Finally, we also measured the pixel-based representation of the images as their “raw” representation.

To measure the distances between representations we computed the Euclidian distance between pairs of faces (python’s numpy linalg.norm method). To compare between dissimilarity measures across different layers and between DCNNs and humans we normalized the distance scores by dividing the measured distances by the maximal distance value in each layer for each of the conditions.

#### Quantifying view-invariance of face-representations in DCNNs

We used two measures to quantify view-invariant face representation:

##### Distances

We computed the Euclidian distances between the representations of the following pairs of faces for the 15 different identities: Same identity faces- same view, Same identity faces – quarter view, Same identity faces – half view (11 pairs), Different identity faces – same view (see Figure 2A). These distances were computed across the different layers of the face-trained, and object-trained DCNNs.

##### Correlations

We computed the Euclidian distances between each identity and all the other identities within the same view condition. This resulted in 15 lists of distances for the reference-view, frontal-view and quarter-left-view conditions (one list per identity), and 11 lists for the half-left-view condition. We then measured the Pearson correlations between these lists of distance scores across the different views (see also (25)). We examined the relative magnitude of the correlations across the different head views for the 15 identities. Since for the half-left distances we only had 11 pairs, we correlated them with the corresponding 11 identities in the reference-view pairs.

#### Quantifying view-invariance of face-representation in humans

To compare the face-representations in DCNNs to that of humans, we used the same correlation-based approach described in the previous paragraph, only now, instead of using distances between representations that were generated by DCNNs, we collected image similarity ranking from human subjects. Each subject rated the similarity between all 105 pairs of images within each of the 4 view-conditions. Similarity was measured on a scale of 1 (very different) to 6 (very similar). In total, 41 participants rated face similarities of pairs of different identities. Each participant was presented with only one of the four head-view conditions (Figure 3A): Reference faces (n= 10), Frontal faces (n= 11), Quarter-left faces (n= 10), Half-left faces (n= 10). Similarity ratings of each image pair within each head view was averaged across participants. We then correlated the similarity ratings of the frontal view, quarter left view and half-left view, with similarity ratings of the reference image pairs across the same pairs of different identities.

### Study 2

#### Stimuli

25 faces were used to generate image pairs. For each of the 25 faces we had an original image, an image in which we replaced critical features (modified from the original image), and an image in which we replaced non-critical features (also modified from the same original image) (for more information about the creation of the face images, see (8)). In addition, we had a different not-modified image of that person, which we used as a reference image. Thus, we created 4 image pairs: 1 – Same pair – the reference image vs. the original image, 2 – Different pair – the reference image vs. a reference image of a different identity, 3 – Critical features pair – the reference image vs. the original image with different critical features, and 4 – Non-critical feature pair – the reference image vs. the original image with different non-critical features (See Figure 4A for example image pairs).

#### Measuring image similarity

We used the same face-trained and object-trained DCNNs as in Study 1, and the same method for generating face representations from the face images. The Euclidian distances between image representations were used to measure image similarity between each pair of images of the 4 conditions described above.

## References

1. N. Kanwisher, Domain specificity in face perception. Nat. Neurosci. 3, 759–763 (2000).

2. D. Y. Tsao, M. S. Livingstone, Mechanisms of face perception. Annu. Rev. Neurosci. 31, 411 (2008).

3. N. Kanwisher, G. Yovel, The fusiform face area: a cortical region specialized for the perception of faces. Philos. Trans. R. Soc. B Biol. Sci. 361, 2109–2128 (2006).

4. P. J. Phillips, et al., Face recognition accuracy of forensic examiners, superrecognizers, and face recognition algorithms. Proc. Natl. Acad. Sci. U. S. A. 115, 6171–6176 (2018).

5. Y. Lecun, Y. Bengio, G. Hinton, Deep learning. Nature 521, 436–444 (2015).

6. R. M. Cichy, D. Kaiser, Deep Neural Networks as Scientific Models. Trends Cogn. Sci. 23, 305–317 (2019).

7. D. L. K. Yamins, et al., Performance-optimized hierarchical models predict neural responses in higher visual cortex. Proc. Natl. Acad. Sci. U. S. A. 111, 8619–24 (2014).

8. N. Abudarham, G. Yovel, Reverse engineering the face space: Discovering the critical features for face identification. J. Vis. 16, 40 (2016).

9. N. Abudarham, G. Yovel, Same critical features are used for identification of familiarized and unfamiliar faces. Vision Res. (2018) https://doi.org/10.1016/j.visres.2018.01.002.

10. N. Abudarham, L. Shkiller, G. Yovel, Critical features for face recognition. Cognition (2019) https://doi.org/10.1016/j.cognition.2018.09.002.

11. N. Kriegeskorte, M. Mur, P. Bandettini, Representational similarity analysis – connecting the branches of systems neuroscience. Front. Syst. Neurosci. 2, 4 (2008).

12. W. A. Freiwald, D. Y. Tsao, Functional compartmentalization and viewpoint generalization within the macaque face-processing system. Science (80-.). (2010) https://doi.org/10.1126/science.1194908.

13. N. Abudarham, G. Yovel, Reverse engineering the face space: Discovering the critical features for face identification. J. Vis. 16 (2016).

14. A. J. E. Kell, D. L. K. Yamins, E. N. Shook, S. V. Norman-Haignere, J. H. McDermott, A Task-Optimized Neural Network Replicates Human Auditory Behavior, Predicts Brain Responses, and Reveals a Cortical Processing Hierarchy. Neuron 98, 630–644.e16 (2018).

15. I. Gauthier, C. A. Nelson, The development of face expertise. Curr. Opin. Neurobiol. 11, 219–224 (2001).

16. I. Gauthier, P. Skudlarski, J. C. Gore, A. W. Anderson, Expertise for cars and birds recruits brain areas involved in face recognition. Nat. Neurosci. 3, 191–197 (2000).

17. H. P. O. de Beeck, C. I. Baker, J. J. DiCarlo, N. G. Kanwisher, Discrimination training alters object representations in human extrastriate cortex. J. Neurosci. 26, 13025–13036 (2006).

18. G. Rhodes, V. Locke, L. Ewing, E. Evangelista, Race Coding and the Other-Race Effect in Face Recognition. Perception 38, 232–241 (2009).

19. M. Wang, W. Deng, J. Hu, X. Tao, Y. Huang, Racial Faces in-the-Wild: Reducing Racial Bias by Information Maximization Adaptation Network (2018) (October 16, 2019).

20. G. Marcus, Deep Learning: A Critical Appraisal (2018) (October 16, 2019).

21. F. Gosselin, P. G. Schyns, Bubbles: a technique to reveal the use of information in recognition tasks. Vision Res. 41, 2261–2271 (2001).

22. M. D. Zeiler, R. Fergus, “Visualizing and Understanding Convolutional Networks” in (Springer, Cham, 2014), pp. 818–833.

23. P. J. Phillips, H. Wechsler, J. Huang, P. J. Rauss, The FERET database and evaluation procedure for face-recognition algorithms. Image Vis. Comput. 16, 295–306 (1998).

24. P. J. Phillips, H. Moon, S. A. Rizvi, P. J. Rauss, The FERET evaluation methodology for face-recognition algorithms. Pattern Anal. Mach. Intell. IEEE Trans. 22, 1090–1104 (2000).

25. I. Blank, G. Yovel, The structure of face-space is tolerant to lighting and viewpoint transformations. J. Vis. 11, 15–15 (2011).

